# Best practice recommendations for sample preservation in metabarcoding studies: a case study on diatom environmental samples

**DOI:** 10.1101/2022.05.04.490577

**Authors:** Baricevic Ana, Chardon Cécile, Kahlert Maria, Karjalainen Satu Maaria, Maric Pfannkuchen Daniela, Pfannkuchen Martin, Rimet Frédéric, Smodlaka Tankovic Mirta, Trobajo Rosa, Vasselon Valentin, Zimmermann Jonas, Bouchez Agnès

## Abstract

The development of DNA metabarcoding and High-Throughput Sequencing for diatoms is nowadays offering an interesting approach to assess their communities in freshwater and marine ecosystems. In the context of the implementation of these genomic methods to environmental monitoring, protocol constraints are moving from scientific to operational applications, requiring operational guidelines and standards. In particular, the first steps of the diatom metabarcoding process, which consist of sampling and storage, have been addressed in various ways in scientific and pilot studies.

The objective of this study was to compare three currently applied preservation protocols through different storage durations (ranging from one day to one year) for phytobenthos and phytoplankton samples intended for diatom DNA metabarcoding analysis. The experimental design included four freshwater and two marine samples from sites of diverse ecological characteristics. The impact of the preservation and storage was assessed through diatom metabarcoding endpoints: DNA quality and quantity, diversity and richness, community composition and ecological index values (for freshwater samples). The yield and quality of extracted DNA only decreased for freshwater phytobenthos samples preserved with ethanol. Diatom diversity was not affected and their taxonomic composition predominantly reflects the site origin. Only rare taxa (below 100 reads) differed among methods and durations. Thus, importance of preservation method choice is important for low-density species (rare, invasive, threatened or toxic species). However, for biomonitoring purposes, freshwater ecological index values were not affected whatever the preservation method and duration considered (including ethanol preservation), reflecting the site ecological status.

This study proved that diatom metabarcoding is robust enough to replace or complement the current approach based on morphotaxonomy, paving the way to new possibilities for biomonitoring. Thus, accompanied by operational standards, the method will be ready to be confidently deployed and prescribed in future regulatory monitoring.

## 1. Introduction

Aquatic ecosystems provide many ecosystem services and functions such as fishing, water provisioning and recreation, and host a wide biodiversity (Grizzetti et al. 2016). However, these ecosystems are subjected to many pressures that can cause physical alteration, water pollution and appearance of invasive species. Facing these pressures, the governmental agencies have implemented monitoring programs of their ecosystems. In Europe, the Water Framework Directive (WFD, European Commission 2000) and the Marine Strategy Framework Directive (MSFD, European Commission 2008) have been set up, respectively, for freshwater and marine ecosystem monitoring. These directives use biological communities to assess the ecological quality of aquatic ecosystems. Diatoms, an abundant group of microalgae, are used as bioindicators in both marine and freshwater ecosystems. In marine ecosystems, plankton diatom communities are present in pelagic areas and some toxic species can bloom. Monitoring of diatom communities enables to follow the dynamics of such toxic species which present a large panel of abundance from rare to highly abundant (AFNOR 2004). In freshwater ecosystems such as rivers and lakes, benthic diatoms are used to assess ecosystems’ ecological quality. Indeed, they present a huge taxonomic diversity (Mann and Vanormelingen 2013) and each species presents particular ecological preferences making them excellent ecological indicators (Rimet 2011). Different biotic indices based on the ecological preferences of the most abundant species have been developed and standardised for the WFD application in European countries (Kelly 2013).

Until now, standard methods used to count and identify diatoms to species level are based on morphological criteria visible by optical microscopy. This method requires time, highly trained taxonomists and can present considerable inter-operator variation (Kahlert et al. 2012). However, during the past decade, the development of DNA metabarcoding and High-Throughput Sequencing offered an interesting alternative (Kermarrec et al. 2013) that can be applied to biomonitoring (e.g. Trobajo et al. 2021). Several studies conducted at regional scale (e.g., 2 cantons in Switzerland with 87 samples: Apothéloz-Perret-Gentil et al. 2017 ; Mayotte Island, France, with 80 samples: Vasselon et al. 2017a ; Catalonia, Spain, with 160 samples: Pérez-Burillo et al. 2021), at national scale (e.g. France with 450 samples: Rivera et al. 2020) and at transnational scale (e.g. in Fennoscandia: Bailet et al. 2019 ; along Danube river with JDS4: Zimmermann et al. 2021) confirmed that this approach is applicable for freshwater monitoring. This was also recently applied to coastal waters (Pérez-Burillo et al. 2022). Marine plankton diversity has been characterised using metabarcoding within important European initiatives like Tara Oceans (de Vargas et al. 2015) and Biomarks (Massana et al. 2015).

The diatom metabarcoding process is composed of five steps: 1) sampling and storage, 2) DNA extraction, 3) PCR amplification, 4) library preparation and HTS sequencing, and 5) bioinformatics treatment. All these steps can show variations among studies with a large range of protocols that are used. To date, only a few studies compared protocols; Vasselon et al. (2017b) compared different DNA extraction protocols; Kermarrec et al. (2013, 2014) compared different barcodes; Vasselon et al. (2018) showed the effect of a cell biovolume correction factor to avoid the quantification bias in metabarcoding; and Tapolczai et al. (2019), Rivera et al. (2020) and Bailet et al. (2020) compared the impact of different bioinformatics pipelines on biotic indices. None of these studies dealt with the first steps of the metabarcoding process, i.e. sampling and storage.

Standardisation efforts at the CEN (European Committee for Standardisation) have already accompanied the application of the European Directives. However, standardization of genomic methods for biomonitoring is still in its infancy and CEN edited in 2018 two technical reports: one about the management of diatom DNA barcodes (CEN 2018a) and one updating the sampling protocol to enable DNA extraction from the samples (CEN 2018b). However, the preservation methods currently used for morphological analyses are based on Lugol’s iodine or formaldehyde, which do not preserve DNA adequately for further DNA-based applications. Alternative preservation methods have been used in various scientific metabarcoding studies: deep-freezing (Visco et al. 2015), ethanol (final concentration 70%) (Vasselon et al. 2017a, Rimet et al. 2018), commercial or homemade preservative solutions (Kelly et al. 2018). The CEN Technical Report published in 2018 (CEN 2018b), is proposing this variety of preservation methods as DNA-friendly alternatives, based on all these studies. However, little is known about the relative effectiveness of these various protocols in sustainably preserving the raw samples. Moreover, to our knowledge, no guidance is available on how long preserved samples should be kept to ensure reliable results using DNA metabarcoding.

In the context of the implementation of genomic methods to environmental monitoring, protocol constraints are moving from scientific to operational applications. The choice of a preservation method by end-users will also have to depend on sampling and shipment operational constraints. For example, during a field sampling day, including the visit of several sites, deep-freezing may be complicated to implement. If a shipment is required, it is easier, cheaper and safer to use a preservative free of hazardous compounds. Moreover, while 100 to 200 samples could be processed per sequencing run, the delay to gather all these samples from the field can take a few weeks to months. So, it is important to know if the preservation protocol and/or the preservation duration have an impact on the final assessment of the diatom community structure, in order to derive best-practices and even standards.

The aim of our study is to highlight best practices for preserving phytobenthos and phytoplankton samples for DNA-based applications in the framework of diatom biomonitoring purposes. To that aim we compared different preservation methods and durations. The recommendations obtained on this basis will be useful to consider in the context of standardization.

We compared three preservation protocols through different storage durations. These three preservation protocols are based on those proposed in the CEN Technical Report on phytobenthos sampling and preservation (CEN 2018b) and described in the literature: 1) ethanol, 2) deep freezing, or 3) nucleic acid preservative. The effect of storage duration was evaluated for the 3 different preservation protocols through 6 different storage durations, ranging from one day to one year. Methods were tested on phytoplankton samples from two marine sites, and on phytobenthos samples from four contrasting river sites in Europe. Preservation methods were compared over time based on 1) the quantity and quality of extracted DNA and 2) the diatom community diversity and structure assessed through DNA metabarcoding (Vasselon et al. 2017a).

## 2. Material and Methods

The experimental design to compare preservation protocols across different storage durations for phytobenthos and phytoplankton samples is summarized in **Figure 1** and detailed on **Supp. Data 1**.

**Figure 1.**
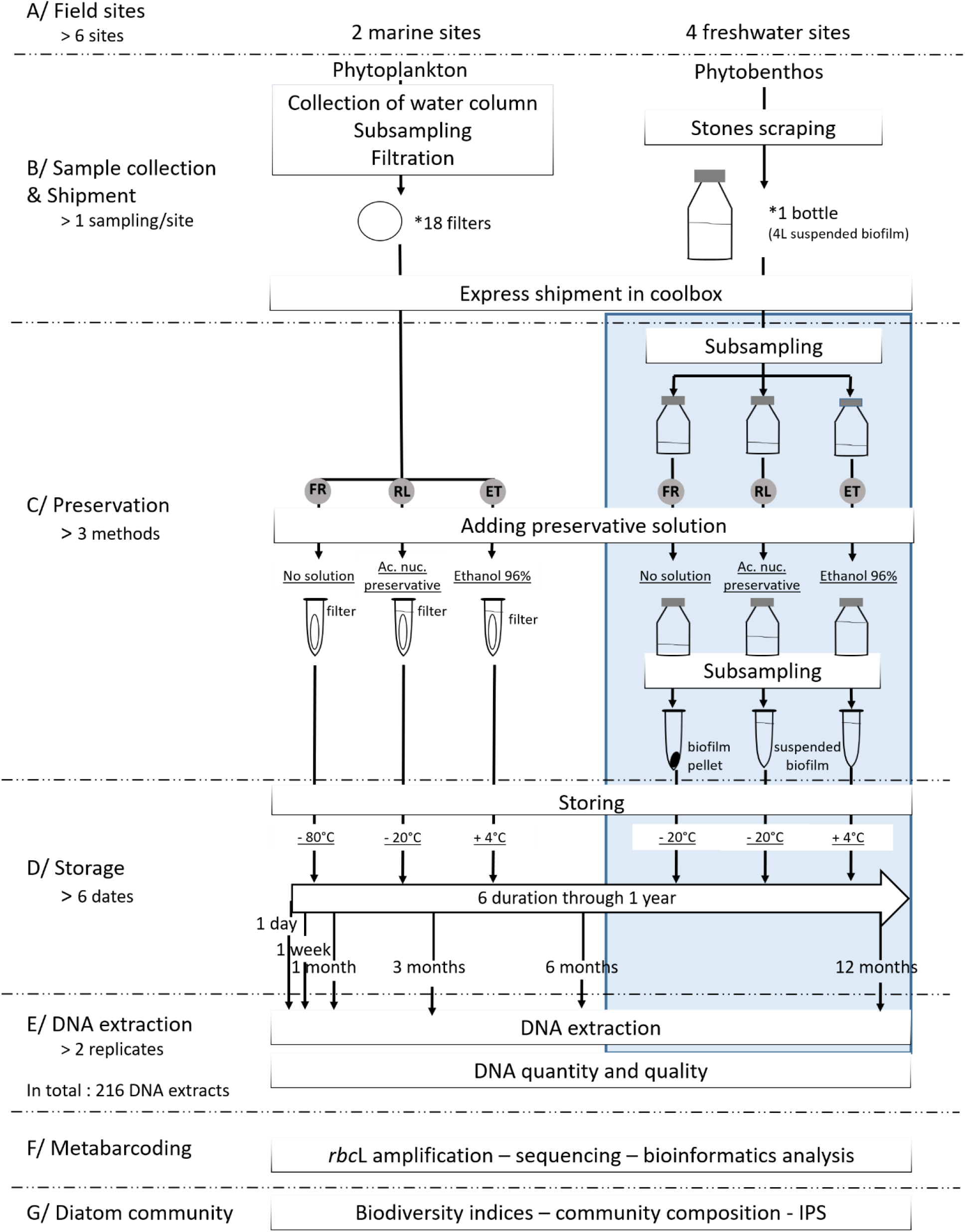
Workflow of the study presenting the three preservation methods (FR: deep-frozen, RL: nucleic acid preservative solution, and ET: ethanol). The blue box is detailed in Supp. Data 1. See Material and Methods for detailed explanations.

### 2.1 Site selection and sampling

Six contrasting European sites (2 Mediterranean marine sites – Spain, Croatia; 4 European river sites – France, Spain, Germany, Finland) were selected for sampling, based on variation in water quality and typology (**Table 1**). Sampling was done on the same day (September 18^th^ 2017) on all selected sites.

**Table 1.** Description of the sampling sites: site code, location, geographic references according to WGS84 system, trophic state, aquatic ecosystem and biotic compartment are indicated.

| Site Code | Location | Latitude | Longitude | Trophic state | Aquatic ecosystem | Biotic compartment |
| --- | --- | --- | --- | --- | --- | --- |
| LC | Lim bay – Croatia | 45,132529 | 13,66059 | mesotrophic | marine | phytoplankton |
| ES | Ebro bay – Spain | 40,816710 | 0,73077 | mesotrophic | marine | phytoplankton |
| OF | Edian river – France | 46,255750 | 6,72342 | oligotrophic | freshwater | phytobenthos |
| MS | Ebro river – Spain | 40,815005 | 0,51997 | mesotrophic | freshwater | phytobenthos |
| EG | Teltow channel – Germany | 52,437615 | 13,32039 | eutrophic | freshwater | phytobenthos |
| HF | Kalimenoja river – Finland | 65,169722 | 25,86889 | humic | freshwater | phytobenthos |

Freshwater phytobenthos was sampled from biofilms following the European standards (AFNOR 2003, 2004) at each of the four river sites (OF, MS, EG, HF). The resulting biofilm suspension was transferred to a sterilized bottle that was stored in a cool box with ice packs for express shipment within 1 day to INRAE lab (Thonon, France) (**Figure 1**). After the 4 river samples had been received, they were sedimented for a few hours and concentrated by removing water supernatant to a final volume of around 900 mL of suspended biofilm per sample. Samples were then homogenised and subsampled into 3 sterilized bottles, one bottle of 300 mL per preservation method.

Marine phytoplankton was sampled by one vertical net haul on both marine sites (LC and ES) with a phytoplankton net (50 μm mesh size) from 15 m deep to the surface. Each net sample was suspended and evenly filtered until complete filter saturation (30 ml per filter for station LC and 60 ml per filter for station ES), on 1.2 μm cellulose (Millipore) (LC site) or GF/F glass microfiber filters (Whatman) (ES site). For each site, 18 filters were obtained in total (6 filters per preservation method) and were placed in marked tubes and stored in a cool box with ice packs for express shipment within 1 day to Center for Marine Research (CMR) lab (Rovinj, Croatia) (**Figure 1**). Samples were subsampled with 2 half-filters as replicates, resulting in 12 half-filters per preservation method.

### 2.2 Sample preservation methods

Three preservation methods were applied to phytoplankton and phytobenthos samples (**Table 2, Figure 1**): 1) deep-frozen (hereafter “FR”) (Visco et al. 2015), 2) preservation with a nucleic homemade acid preservative solution (hereafter “RL”, standing for “RNA Later” style) (Kelly et al. 2018) and 3) preservation with ethanol (hereafter “ET”) (Vasselon et al. 2017a).

**Table 2.** Description of the three preservation methods: method code, biotic compartment, storage conditions and material used for extraction are indicated.

| Preservation method name | Preservation method code | Biotic compartment | Fixative solution | Storage temperature | Material used for preservation | Material used for extraction |
| --- | --- | --- | --- | --- | --- | --- |
| Cryopreservation | FR | Phytobenthos | no | -20°C | Pellet | Pellet |
|  |  | Phytoplankton | no | -80°C | Filter | Filter |
| DNA stabilization solution preservation | RL | Phytobenthos | home-made nucleic acid preservative | -20°C | Suspended biofilm with fixative solution | Pellet |
|  |  | Phytoplankton | home-made nucleic acid preservative | -20°C | Filter with fixative solution | Filter |
| Ethanol preservation | ET | Phytobenthos | Ethanol (final conc.~70%) | +4°C | Suspended biofilm with fixative solution | Pellet |
|  |  | Phytoplankton | Ethanol (final conc.~96%) | +4°C | Filter with fixative solution | Filter |

### FR preservation method

For freshwater samples, 12 2mL-subsamples of the biofilm suspension were obtained from one 300mL bottle under agitation for each site. Subsamples were then centrifuged and pellets were frozen and stored at -20°C (**Supp. Data 1**). For marine samples, 12 half-filters were frozen and stored in tubes at -80°C for each site.

### RL preservation method

A nucleic acid preservation solution was home-made with 3.5 M ammonium sulphate, 17 mM sodium citrate and 13 mM Ethylene-diamine-tetraacetic acid (EDTA). pH was adjusted to 5.2 using 1M H_2_SO_4_ and solution was sterilized by filtration with 0.2 μm filter. For freshwater samples, one volume of the nucleic acid preservative solution was added to one volume of sampled biofilm, for one 300mL bottle under agitation. Four 2mL-subsamples of the preserved biofilm suspension were then stored for each site (**Supp. Data 1**). For marine samples, 2 mL of the preservative solution was added in 12 tubes with half-filters for each site. All samples were then stored at -20°C.

### ET preservation method

For freshwater samples, three volumes of 96% ethanol were added to one volume of biofilm, in order to obtain a final ethanol concentration above 70%. This was applied to one 300mL bottle under agitation. A 17mL-subsample of the preserved biofilm suspension was then stored for each site (**Supp. Data 1**). For marine samples, 2 mL of 96% ethanol was added in 12 tubes with half-filters. All samples were then stored in the dark at +4°C.

In all subsampling phases for freshwater biofilm samples, special attention was paid to homogenisation by 1) permanent agitation of the solution to subsample and 2) sequential subsampling, adding solution to all subsamples in succession, 1mL per 1mL. Marine net samples were also well homogenised during filtration procedure to enable even distribution of the sample on each filter.

### 2.3 Preservation duration and DNA extraction

The samples, preserved with the three methods, were further processed at six different preservation durations (1 day, 1 week, 1 month, 3 months, 6 months and 1 year) during one year (**Figure 1**). For each duration, 2 replicates were retrieved per preservation method and per site (i.e. 36 samples) and independently processed for DNA extraction. For marine samples, replicates were obtained by cutting each filter in 2 halves independently processed for DNA extraction.

For freshwater samples, DNA extraction was performed on biofilm pellets, either those directly preserved (FR samples, **Table 2, Supp. Data 1**) or those obtained by centrifugation just before DNA extraction (RL and ET samples, **Table 2, Figure 1**). In order to minimize the dilution effect of RL and ET methods (respectively ½ and ¼), DNA extraction was processed on the same total amount of biofilm 2mL-pellets for each preservation method (1 pellet for FR, 2 pooled pellets for RL and 4 pooled pellets for ET). For marine samples, half-filters were used directly for DNA extraction.

DNA extractions were performed using a commercial kit (Macherey–Nagel NucleoSpin® Soil kit, Duren Germany) with purification columns following Vautier et al. (2020). In short, biofilm pellets and phytoplankton filters were resuspended in lysis buffer and mechanically disrupted using ceramic beads. After proteins and undissolved sample material precipitation, supernatant with dissolved DNA was first passed through inhibitor removal columns and next through NucleoSpin® Soil columns for DNA binding where PCR inhibitors were removed by efficient washing. Finally, DNA was eluted in the NucleoSpin® Soil elution buffer (Tris/HCl buffer) and stored on -20°C prior PCR amplification and High-Throughput Sequencing (**Figure 1**). DNA extractions were processed at INRAE lab (Thonon, France) for freshwater samples, and at CMR lab (Rovinj, Croatia) for marine samples. All DNA extracts from marine samples were then sent with express shipment to INRAE lab for downstream analysis at the end of the 1-year period. In total, for this 1-year experiment, 216 DNA extracts were obtained.

### 2.4 DNA quality and quantity

At the end of the 1-year preservation period, DNA quality and quantity were assessed on all 216 DNA extracts (**Figure 1**). To evaluate the DNA quality, the 260/280 nm ratio was measured by spectrophotometry with Nanodrop®ND-1000 (Nanodrop Technologies, Wilmington, Delaware). To evaluate the DNA quantity, DNA concentration (ng/μL) was measured on two replicates per DNA extract with the Life Technologies (Carlsbad, California) Quant-iTTM PicoGreen® dsDNA assay kit using a microplate reader (Fluoroskan AscentTM FL; Thermo Scientific, Waltham, Massachusetts) and following manufacturer’s instructions. Mean concentration value of the sample replicates was used in analyses.

### 2.5 PCR amplification and sequencing

As proposed by Vasselon et al. (Vasselon et al. 2017a), a 312 bp fragment of the *rbc*L chloroplastic gene was amplified from DNA extracts using Takara LA Taq® polymerase and an equimolar mix of the forward primers Diat_rbcL_708F_1, 708F_2, 708F_3 and the reverse primers R3_1, R3_2, following protocol by Chardon et al. (2020). For each DNA sample, PCR amplification was performed in triplicate in a final volume of 25 μL. 19 μL of each replicate were pooled together and 50 μL of this pool were transferred in an individual well of a 96-well microplate. The resulting 3 plates with 216 samples were sent to “GenoToul Genomics and Transcriptomics” (GeT-PlaGe, Auzeville, France) where library preparation, final library pool and the sequencing with Illumina MiSeq System with paired-end sequencing kit (V2, 250 bp × 2) were performed (**Figure 1**).

### 2.6 Bioinformatics

Demultiplexing was performed by the sequencing service and two fastq file (forward and reverse) were provided for each library prepared (2 × 216 fastq files in total). Fastq files were processed using the Mothur software (Schloss et al. 2009). Contigs were made by merging forward and reverse reads trimmed to only the overlapping section. Further, reads in every fastq file were quality filtered by excluding sequences with overlap shorter than 180 bp and all sequences with ambiguities and mismatches. From the resulting good quality reads exact duplicates were removed (dereplication). Next bioinformatics steps on good quality reads processing included alignment of these sequences to the reference alignment and preclustering that enabled de-noising of sequences (1 difference for every 100 bp of sequence was allowed). Chimera removal was done using the VSEARCH algorithm with default parameters. Sequence classification was made using the naïve Bayesian Method (Wang et al. 2007) with bootstrap confidence score set to 85% and Diat.barcode v7 as reference library (Rimet et al. 2016, 2019). Sequences classified to taxa other than diatoms (Bacillariophyta) were removed from further processing. Operational taxonomic units (OTUs) were defined by calculating distances between sequences and clustering these distances based on 0.05 difference cut-off implementing the furthest neighbour algorithm. OTUs containing one single sequence (singletons) were removed. Sample replicates were merged (merge.groups) and replicate reads were summed to get read abundances. Consensus taxonomy for each OTU was assigned at 85% confidence threshold using reads taxonomy as introduced by the classify.otu command. A list of taxa and their relative abundances (based on read abundances) in each sample was produced. OTUs with identical highest taxonomy level were merged (merge.otus). For normalizing the data, random subsampling was performed with size (number of reads) set to the size of the smallest sample.

### 2.7 Statistical analyses

Statistical analyses as well as graphical presentations of the results were performed using the R software and included core packages (R Core Team 2017). To find statistically significant differences (p<0.05) in DNA concentration and diatom community diversity among different preservation duration and preservation methods, two-way ANOVA was used. Spearman’s rho statistic was used to estimate a rank-based measure of association in correlation analyses. Patterns of sample dissimilarity were visualized using unconstrained ordinations of non-metric multidimensional scaling (nMDS). Ordinations were based on taxa presence/absence and abundance using Bray-Curtis index. Further, to find statistically significant differences (p<0.05) in the detected diatom community compositions, permutational multivariate ANOVA (PERMANOVA) was used with the adonis function in vegan (Oksanen et al. 2018) with number of permutations: 9999. Community composition was also compared by Venn diagrams using vegan package in R. Dominant and rare taxa presence/absence for different preservation durations and methods were analysed for each sampling station.

### 2.8 Diatom indices calculation

For freshwater river sites, we assessed their ecological quality using the Specific Pollution-sensitivity Index (SPI) (Cemagref 1982) based on species inventories (species composition and relative abundances based on read numbers; or genus, if species level was not reached) obtained by metabarcoding. SPI values were calculated using the OMNIDIA 5 software (Lecointe et al. 1993).

## 2.9 Data availability

>> address for Fastq files in open access on data.inrae.fr with DOI (to be added when manuscript is ready for publication)

## 3. Results

### 3.1 DNA quality and quantity

Spectrophotometry measurements confirmed good DNA quality with 260/280 nm ratios between 1.8 and 2 for all samples. Measured DNA concentrations differed among samples and ranged from 1 to 160 ng/μL (**Figure 2**) with marine samples having significantly lower concentrations (mean value 7.1 ng/μL) than the freshwater ones (mean value 65.8 ng/μL). Effect of the preservation time and method was analysed on the DNA concentrations. The ANOVA analysis revealed that preservation methods had an effect on DNA concentration (p < 0.001) with ET samples differing significantly from FR and RL samples. This effect of ethanol preservation on DNA concentration was present only for freshwater samples for which ET samples had significantly lower DNA concentrations (p < 0.001) compared to the other two types of preserved samples (FR, RL). Preservation duration did not have an effect on DNA concentration (p >> 0.05) for FR and RL methods. However, for ET method a significant decrease in DNA concentration was observed for freshwater samples that was particularly marked between 1- and 3-months durations.

**Figure 2.**
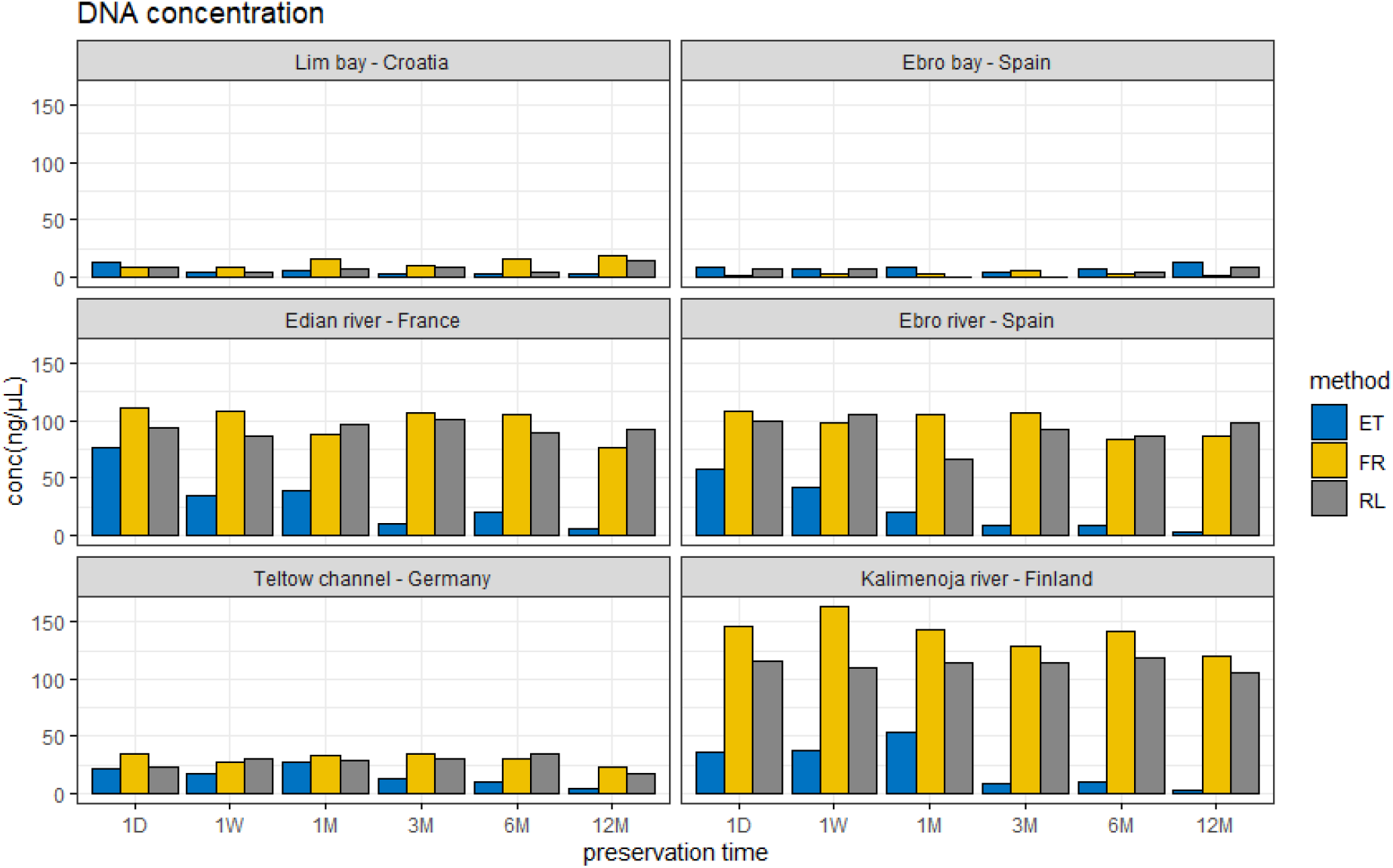
DNA concentrations (mean values of 3 replicates) over time (x axis) for the three preservation methods (blue: ET, yellow: FR, grey: RL) for marine sites (top row) and freshwater sites (middle and last rows).

### 3.2 Diatom community diversity

High-throughput sequencing of the 216 samples resulted in 2 × 216 fastq files with a total of 7.9 million reads. After bioinformatics processing, a total of 3.9 million reads were conserved with an average of 36621 reads per sample. Reads were clustered in an average number of 97 OTUs/sample and since OTUs with identical highest taxonomy level were merged during processing, the final number of OTUs corresponds to the total number of different diatom taxa detected in the experiment. Rarefaction curves for each sample (**Supp. Data 2**) indicate sufficient sequencing effort has been made to get a reasonable estimate of the taxa number. Sample replicates from the same sample over all methods and durations reveal a significant correlation between reads (R=0.71, p<2.2e-16) and OTU number (R=0.93, p<2.2e-16). Homogeneity of numbers of reads and OTUs between replicates belonging to the same sample confirm the good quality and validity of our dataset.

When all methods and durations are considered, freshwater sites are on average characterised with higher number of reads (41222 reads/sample), higher OTU richness (111 OTUs/sample) and higher diversity index (Shannon) values compared to marine sites (27155 reads/sample and 67 OTUs/sample) (**Figure 3**). The average number of OTUs over all sites is very similar for all three different preservation methods. For each site, preservation methods have no significant impact (one-way ANOVA, p>0.05) on read numbers, OTU richness and Shannon index values (**Figure 3**). For each site, these diversity parameters did not change significantly (one-way ANOVA p>0.05) over time as well.

**Figure 3.**
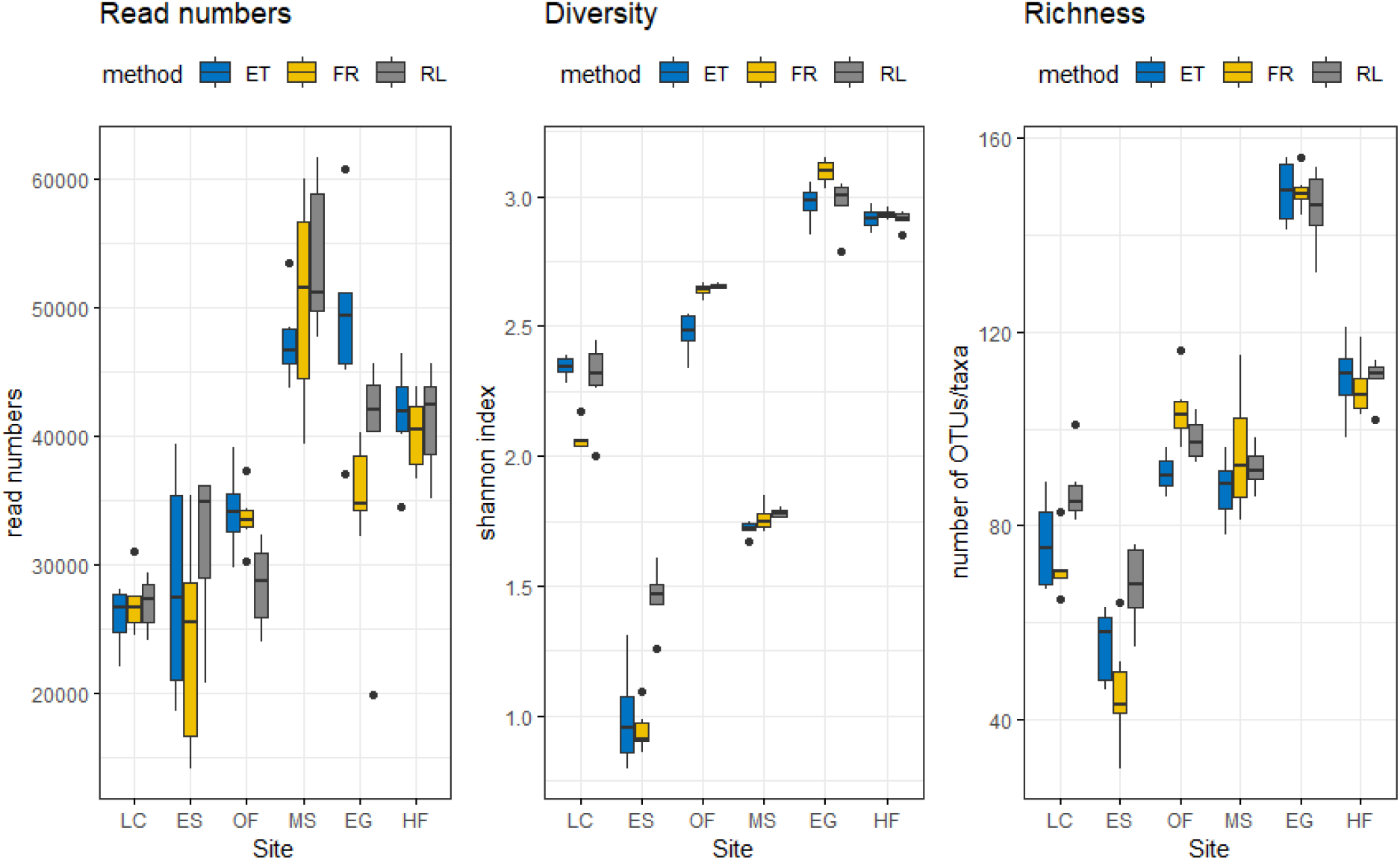
Diversity parameters of diatom communities: box plots of Read numbers, Shannon index and OTU numbers (Richness) for the three preservation methods (blue: ET, yellow: FR, grey: RL) for marine sites (left) and freshwater sites (right).

### 3.3 Diatom community composition

Read subsampling was randomly performed to 14190 reads per sample, based on the sample with the lower number of reads (ES marine sample). OTUs were taxonomically assigned at species or genera level using the Diat.barcode reference library, corresponding to 102 different diatom genera. The diatom community composition differs among sites (**Supp. Data 3**), reflecting their environmental and geographical characteristics (e.g., freshwater vs. marine; oligotrophic vs. eutrophic), and consistent with the known composition of the community for the study year period (comm. pers.). The genus *Nitzschia* was the most abundant in terms of read number and included the highest number of species (28 different species). About 50% of the 102 genera were represented by a single species in the dataset.

#### 3.3.1 Do preservation methods and/or duration impact community structure?

Community ordination taking read abundance into account (diversity, Bray-Curtis metrics) showed (**Figure 5**) that the samples were discriminated according to sampling sites primarily (PERMANOVA, R2= 0.96, p<0.001) and preservation methods (PERMANOVA, R2= 0.01, p<0.001), but not according conservation time (PERMANOVA, R2= 0.0008, p>0.05). Taking presence/absence (richness, Jaccard metrics) into account also confirms sampling sites (PERMANOVA, R2= 0.90, p<0.001) and methods (PERMANOVA, R2= 0.01, p<0.001) significantly influenced the community structure, and time did not (PERMANOVA, R2= 0.002, p>0.05). Site by site analysis showed that preservation methods explained on average 66 % of the total variance in distance between samples, while preservation durations explained on average 11% (**Table 3**). However, looking at community composition at each site through methods and time, we can see that these changes are mainly due to changes in relative abundances for abundant taxa (**Supp. Data 3**) and to changes in presence-absence for low abundant taxa (not shown).

**Figure 5.**
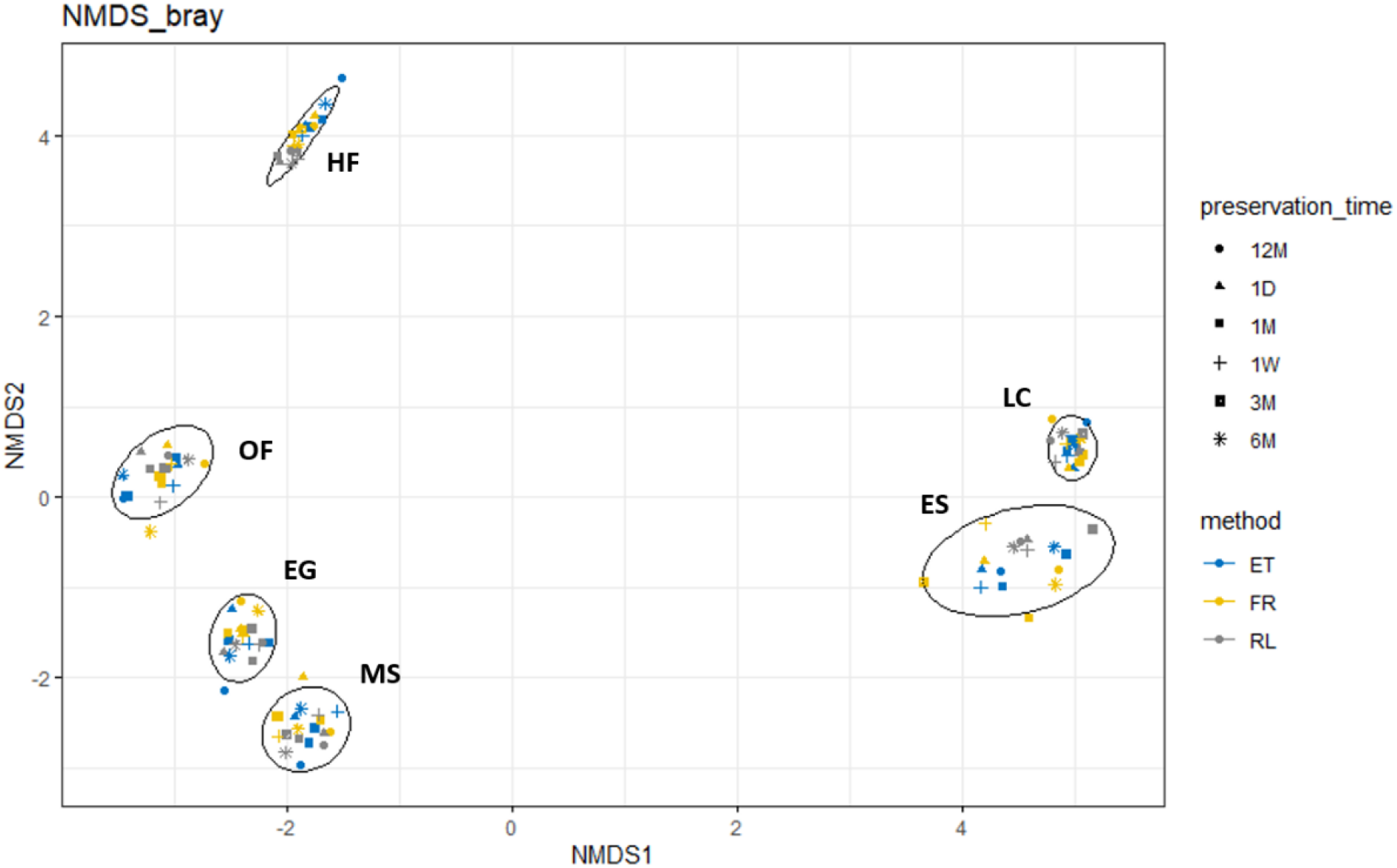
Non-metric multidimensional scaling (NMDS) ordination based on Bray-Curtis distances, taking read abundance into account. Samples are marked according to preservation method (colour) and duration (shape). The 6 sampling sites (HF, OF, EG, MS, ES, LC) are visualised by ellipses.

**Table 3.** Results of PERMANOVA analysis (adonis function) of OTUs, indicating the percentage of variance (R2) explained by preservation method and duration, and associated probability (p).

| site | preservation method |  | preservation duration |  |
| --- | --- | --- | --- | --- |
| | $R^2$ (%) | $p$ | $R^2$ (%) | $p$ |
| LC | 0.546 | 0.0001 | 0.181 | 0.25 |
| ES | 0.574 | 0.0011 | 0.116 | 0.75 |
| OF | 0.605 | 0.0001 | 0.149 | 0.37 |
| MS | 0.726 | 0.0002 | 0.082 | 0.57 |
| HF | 0.849 | 0.0001 | 0.047 | 0.54 |
| EG | 0.683 | 0.0001 | 0.105 | 0.47 |

#### 3.3.2 Are some taxa differentially detected?

There is no significant difference in the number of taxa detected between the preservation methods and the overall number of taxa detected for each method is around 300 taxa/sample. In the dataset, 81% of detected taxa were shared between the three methods, while each method is characterised with less than 5% of taxa detected only by that single method (ET: 11 taxa, RL: 13 taxa, FR: 5 taxa, **Figure 6**). This part of the community corresponds to rare taxa (less than 100 reads, **Figure 6**). Taxa that are unique to one method are taxonomically diverse and do not share evident ecological characteristics. The rare community also contributes to methods shared-community (36%, **Figure 6**). When looking at the communities according to preservation durations, 75% of the diatom community is shared, while communities unique to each duration corresponds to low abundant taxa (less than 100 reads). Only one non-rare species was unique to first duration (1 day) and RL method: *Actinoptychus splendens*, which is a marine species detected only at the ES site (Ebro bay, Spain). Rare taxa are often method-specific and usually appear and disappear over time without any obvious pattern.

**Figure 6.**
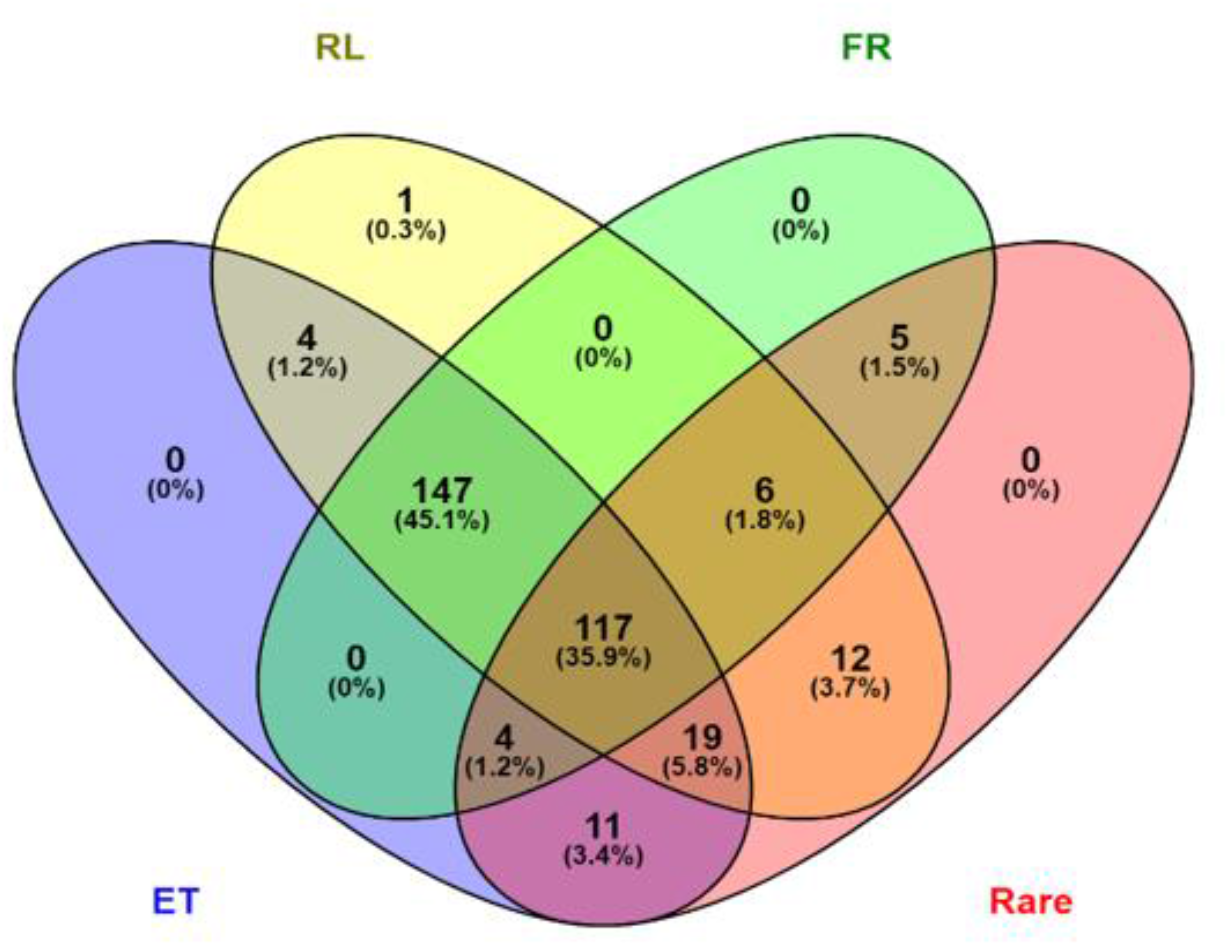
Comparison of the number of shared diatom taxa between the three preservation methods (ET: blue, RL: yellow, and FR: green). Rare taxa (less than 100 reads) are presented in red.

### 3.4 Ecological quality index for freshwater sites

Based on OTUs assigned at species or genus levels and their read abundances, SPI scores were calculated for freshwater sites. They ranged from 14.2 - 18.9 (**Figure 7**). Eutrophic (EG) and humic (HF) freshwater sites had lower mean SPI values, 14.52 and 14.90 respectively. Mesotrophic (MS) and oligotrophic (OF) sites had higher mean SPI values, 17.51 and 18.23 respectively. The influence of sampling site on SPI values was significant (one-way ANOVA, p <2e-16). Most importantly, at each site, SPI values are very stable whatever the preservation method and the duration (no significant change, one-way ANOVA p>0.05).

**Figure 7.**
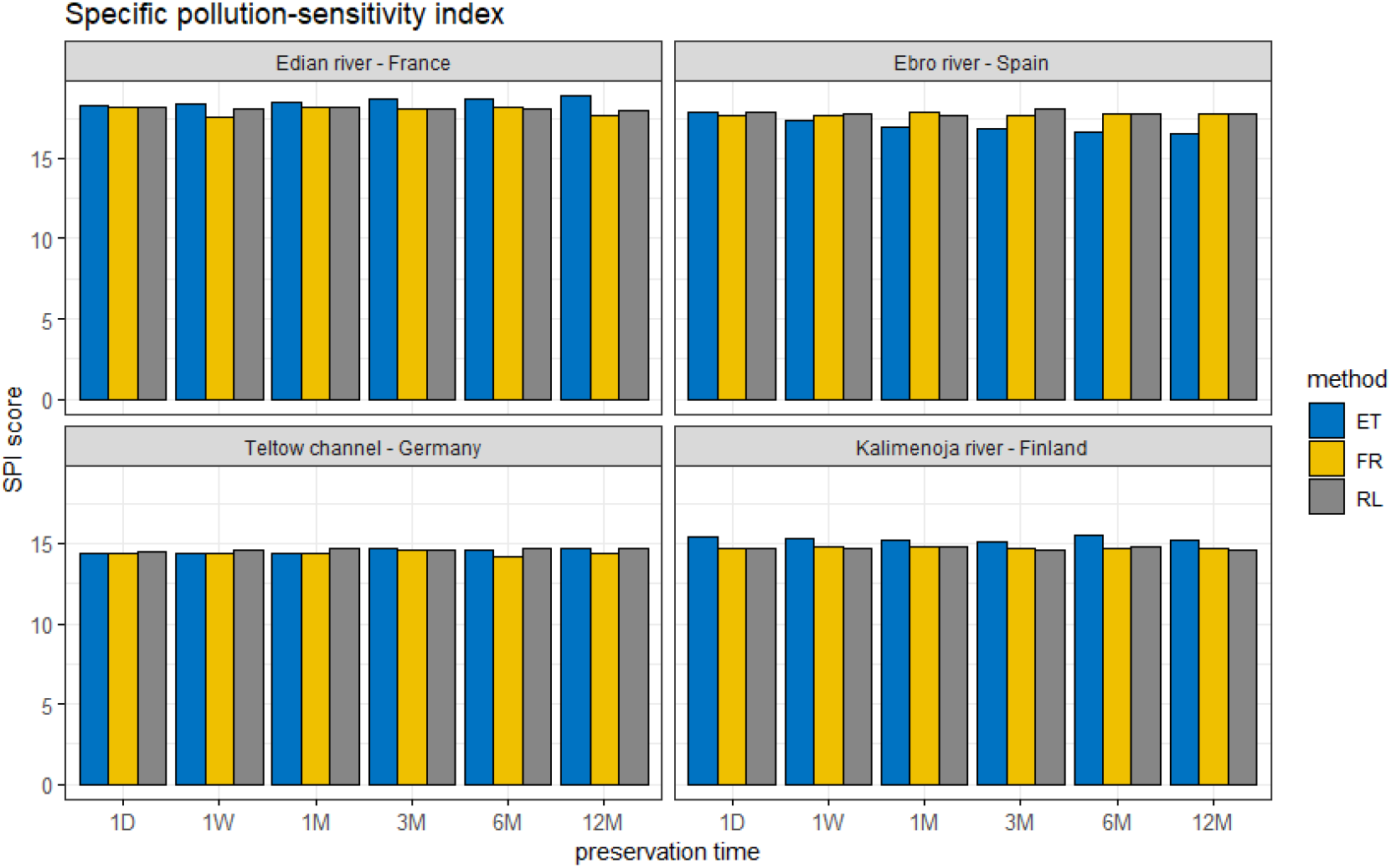
SPI (Specific pollution-sensitivity index) index values over the 3 preservation methods (ET: blue, FR: yellow, RL: grey) and the 6 durations (x axis).

## 4. Discussion

Identification of diatom communities in environmental samples through DNA metabarcoding is nowadays becoming a reliable approach that has been successfully tested on a large variety of ecological contexts through numerous pilot studies for freshwater biomonitoring (Vasselon et al. 2019, Rivera et al. 2020, Pissaridou et al. 2021, Tapolczai et al. 2021, Pérez-Burillo et al. 2020), while less has been done for phytoplankton in marine ecosystems (Piredda et al. 2018, 2022). With sample preservation and storage being at the beginning of this metabarcoding approach, this experiment aimed at identifying potential key points in preservation methods and preservation duration that may blur the ability of DNA metabarcoding to produce accurate diatoms inventories and, as a result, produce reliable quality assessment of aquatic ecosystems.

### 4.1. Sample preservation: a robust first step in diatom metabarcoding

Our results show an overall outstanding robustness of the approach that is seldom affected by the method used to preserve the samples or by the storage duration. Overall, diatom community composition differed among sampling sites and not because of preservation methods neither storage duration. Detecting an important impact of sampling site on community composition is not surprising since sampling was conducted to represent very diverse environments characterised with various freshwater and marine locations as well as various trophic status. Diatoms are known to have specific ecological preferences thus making their communities shaped by sites ecology. This is the reason why these communities are used as Biological Quality Elements (BQEs) to monitor the ecological status of waterbodies in the WFD (Rimet 2011). Very little impact has been observed from the preservation methods and the storage duration on the final endpoints delivered by metabarcoding: community composition, diversity indices and SPI values. Although the water chemistry was different from one site to another, this did not have a differential impact between preservation methods. In all cases the site effect appears to be major and well reflected. The longest storage duration tested here was 1 year after sampling. However, as no significant changes have been observed throughout this period for any endpoints, it may well indicate that longer storage may not affect the final results. This was observed by the authors in previous studies where samples stored over a long period of time and later treated with metabarcoding still provided coherent results with the morphotaxonomy results for RL (Kelly et al. 2020) and for ET (Bailet et al. 2019, Kahlert et al. 2021).

In most cases, the preservation methods we explored do not affect the quantity and quality of the DNA extracted from preserved samples. The exception is the ET method applied to freshwater samples. Preservation with ethanol seems to lead to lower DNA concentrations than other methods. Ethanol acts as both a killing and a preservative agent, replacing water molecules in biological tissue (Carter 2003), and has been successfully used for macroinvertebrates specimen preservation for biomonitoring (Stein et al. 2013). A minimum of 70% ethanol is necessary to ensure the fixation of the samples, preventing their evolution through time due to biotic processes. To attain that, 1 volume of suspended biofilms needs the addition of 3 volumes of 96% ethanol. For the 2 other methods, dilution was reduced (1/2 for RL method) or absent (FR method). However, this higher dilution in ethanol (1/4), compared to RL and FR methods, was compensated by more material (4 biofilm pellets) for DNA extraction, instead of 2 or 1, for RL and FR respectively. Therefore, different dilutions should not impact on the final performance of the DNA extraction.

Recent studies have shown for macroinvertebrate samples that organismal DNA is released from cells into the ethanol used for preservation during sample storage (Martins et al. 2019, Zizka et al. 2019), confirming the replacement of water molecules by ethanol in biological tissues (Carter 2003). The DNA released in ethanol has been suggested and tested as an efficient method to inventory macroinvertebrate communities (Hajibabaei et al. 2012). In our case, some DNA may have also been released from diatom cells and consequently not included in the centrifuged sample that is used for DNA extraction, thus limiting the initial DNA availability. This may have also been facilitated by the absence of freezing step in ET method. Moreover, samples conserved at 4°C may be more prone to abiotic degradations than frozen ones. Finally, biofilms are complex samples that include many organisms (mainly bacteria, microalgae, fungi) embedded in an exopolysaccharide matrix (Flemming and Wingender 2010) and even hosting eDNA from external organisms (Rivera et al. 2021a, 2021b). The extracted DNA is thus a mix of DNA from different organisms. Differential DNA release may occur during the preservation phase in ethanol, e.g. ethanol is releasing more efficiently DNA from algae cells than from other components of the biofilm, potentially adding one more dilution for diatoms.

Preservation duration, from 1 day to 1 year, does not affect the quantity and quality of the DNA extracted from preserved samples, except when preserving freshwater samples with ethanol. In that case a decrease of DNA concentration was observed that was marked after 1 month of preservation, but this trend did not increase until 1-year preservation. We can hypothesize that this decrease is linked to the release of DNA from the cells to the ethanol solution. This could be evaluated by extracting in parallel DNA from the pellets and DNA from the ethanol.

For all methods and dates, even in the “worst case” of ethanol preservation for the samples that have been stored the longest, the final endpoints are not affected. Indeed, the community composition is predominantly homogeneous in each site, whatever the method and the duration of the storage. The small percentage of taxa that differ from one method to the other or from one date to the other are those that are rare (under 100 reads). When diatom metabarcoding is dedicated to the evaluation of ecological status to compare changes in community structure through time and space, which is currently its main application, it is definitely an approach that is not affected by the sample conservation. If diatom metabarcoding is dedicated to the detection of rare species (invasive alien species, endangered species, toxic species), then it may be affected by sample conservation. Low-density invasive or threatened species as well as toxic species with the potential to form harmful blooms could constitute this rare community and for detection of such species, the preservation method choice could be of even greater importance. However, in our study we could not identify a specific trend and derive best practice for that. For that purpose, a higher sequencing depth is probably required to avoid overlooking rare species and needs to be calibrated appropriately. Moreover, in such cases, specific study design (e.g., biological replicates, positive and negative controls) and biomolecular methods focusing on the target species may be more adapted (e.g. dPCR, qPCR).

### 4.2. Towards a standardised and user-friendly method

Following results from the numerous pilot studies, we can be confident that diatom metabarcoding is robust enough and can replace or complement the current approach based on morphotaxonomy. To do so, stakeholders call for guidelines and/or standards to accompany the deployment of the method for biomonitoring purposes (Blancher et al. 2021). General guidelines have been recently published, providing best-practices for many applications of DNA metabarcoding (Bruce et al. 2021, Pawlowski et al. 2021) which represent an important milestone. However, guidelines need to be even more precise to be easily and reliably handled by the operators in a variety of contexts. Diatom metabarcoding includes several steps which require different expertise (e.g. field sampling, molecular analysis, bioinformatics, ecology). For that reason, a step-by-step standardisation would be necessary to 1) enable the commitment of multiple actors/expertise and 2) leave the door open to future technological changes and evolutions that are numerous in the biomolecular sector.

However, methods have to remain operational and as much as possible user-friendly. Concerning sample conservation, depending on usage, one method or the other maybe more adapted. Methods requiring freezing or deep-freezing conservation (FR and RL) imply immediate storage. They may be operational for scientific purposes; however, they require fast access to -80°C or -20°C frozen facility. For operators that have to handle large field campaigns or to access adverse environments, without access to lab facilities during several days, the more operational process is the addition of a preservative solution (nucleic acid preservative or ethanol) directly in the field. However, for biofilm samples, after addition of the nucleic acid preservative, a pellet has to be obtained by centrifugation prior to -20°C freezing which may be difficult to operate in the field (and additional centrifuge facility for biofilms). In this study, we did not test the impact of the conservation time of biofilms in nucleic acid preservative, prior to centrifugation and -20°C storage, which may be an interesting alternative. Field collections become compromised when sample processing cannot be completed within short critical time periods when necessary capacities are unavailable. However, from Ladell et al. (2019) results we can suppose that the conservation of samples in preservative is little affected by storage temperature (frozen or room temperature) in the first week following sampling, thus providing flexibility in the first sampling and storage step of the metabarcoding process. Finally, preservation with ethanol appears to be the most operational strategy for large or remote field campaigns. Moreover, storage in dark at +4°C is also facilitating the easy storage of a large number of samples. For studies not oriented towards the detection of rare species, but rather towards the identification of diatom communities, our results show that ethanol long-term preservation is well adapted.

A first attempt for standardisation has been done in 2018, with the publication of a technical report (CEN 2018b) in line with an existing standard developed in CEN to accompany the WFD. This technical report concerned the initial sampling phase to make it evolve in a DNA-friendly way, compatible with DNA metabarcoding. Indeed, the sample conservation proposed by the initial standard (e.g. lugol, formaldehyde) was preventing the extraction of DNA from conserved samples. Based on ongoing studies at this time a large choice of already tested DNA-friendly conservation methods has been proposed in a technical report (CEN 2018b). On that basis, the present study was conducted to ascertain and precise the use of these conservation methods. From our results, the initial sampling step of the diatom metabarcoding process appears to be robust whatever the preservation method that is used and whatever the storage duration, up to one year and probably longer. Consequently, preservation method choice in future diatom metabarcoding studies could be primarily based on experimental design, field and lab facilities, shipping constraints and available funding and less on necessary choice of one optimal sample preservation method.

Such robustness has been already observed for other steps of diatom metabarcoding: DNA extraction methods (Vasselon et al. 2017b), PCR amplification (Vasselon et al. 2021) bioinformatic processing (Bailet et al. 2020, Rivera et al. 2020). In all these studies, final endpoints, especially ecological quality indices were seldom affected by changes in methods. An open-access reference library, Diat.barcode (Rimet et al. 2019), and related standards (CEN, Rimet et al. 2021) are also completing the toolbox.

## 5. Conclusion and perspectives

This study has shown that preservation method and storage duration seldom affected the DNA metabarcoding results: yield and quality of extracted DNA, diatom community diversity, ecological quality assessment. The decrease in yield and quality of extracted DNA observed only for freshwater phytobenthos samples preserved with ethanol did not affect the community structure and the final index values. Only low abundant taxa differed among methods and durations. Thus, importance of preservation method choice is important for low-density species (rare, invasive, threatened or toxic species). However, for biomonitoring purposes, freshwater ecological index values were not affected whatever the preservation method and duration considered (including ethanol preservation), well reflecting the site ecological status.

Diatom metabarcoding has shown to be robust enough to replace or complement the current approach based on morphotaxonomy, paving the way to new possibilities for biomonitoring (Keck et al. 2017, Pawlowski et al. 2018, Trobajo et al. 2021). Diatom metabarcoding has proven to be less prone to identification errors, to provide high-throughput intercomparable data, allowing its application at large temporal and spatial scales. Due to the high polymorphism of the *rbc*L barcode that is sequenced, the produced data are providing a more detailed scale of observation, often at intraspecific level (Chonova et al. 2021) allowing to better understand the evolution of diatom communities through time and space and their biogeography. Even more, new developments such as artificial intelligence may provide empowered methods, making full use of the genetic signal (Tapolczai et al. 2019, Feio et al. 2020, Apothéloz-Perret-Gentil et al. 2021).

Thus, accompanied by operational standards, the method will be ready to be confidently deployed and prescribed in future regulatory monitoring. Since 2020, CEN has dedicated one of its working group (EN/TC 230/WG 28 - DNA and eDNA methods) to the development of new standards for genomic approaches applied to the biomonitoring of aquatic ecosystems. The results of this study will facilitate the emergence of a new standard, building on the initial technical report (CEN 2018b) and specify its contours. Indeed, the standard may propose flexibility in methods and storage duration, not affecting the final ecological assessment. However, new interlaboratory ring-tests (e.g. Vasselon et al. 2021) will be required to develop the step-by-step standards for diatom metabarcoding. Thus, accompanied by operational standards, the method will be ready to be confidently deployed and prescribed in future regulatory monitoring.

## Acknowledgments

This work was initiated and supported by the DNAqua-Net COST Action CA15219 ‘Developing new genetic tools for bioassessment of aquatic ecosystems in Europe’ funded by the European Union. DNAqua-Net funded the lab exchanges of the two first-authors A. Baricevic and C. Chardon through Short-Term Scientific Missions in 2017 and 2019, respectively. This work largely benefited from the discussions with all participants to the “diatom workshop” held in Limassol (Cyprus) on October 1-2, 2019, organised by CUT and supported by DNAqua-Net, especially S. Derycke (Belgium), T. Elersek (Slovenia), S. Fazi (Italy), M. Kelly (UK), M. Kelly-Quinn (Ireland), Z. Ljubesic (Croatia), S. Theroux (USA), G. Varbiro (Hungary), M. Vasquez (Cyprus). INRAE funded the DNA sequencing at INRAE Genomics (GeT-PlaGe, Auzeville, France). RT acknowledges support of the CERCA Programme/Generalitat de Catalunya, and help from IRTA technicians (D. Mateu, J.L. Costa and M. Rey) for sampling. JZ acknowledges support by the Federal Ministry of Education and Research [German Barcode of Life 2 Diatoms (GBOL2), grant number 01LI1501E]. MP acknowledges support of the CMR research vessel “Burin” crew and of the Croatian Science Foundation project: Life strategies of phytoplankton in the northern Adriatic (UIP-2014-09-6563). MK acknowledges support by the Swedish Agency for Marine and Water Management. CC, FR, VV and AB acknowledge support by the Office Français de la Biodiversité (OFB).

## Supplementary data

**Supp. Data 1.**
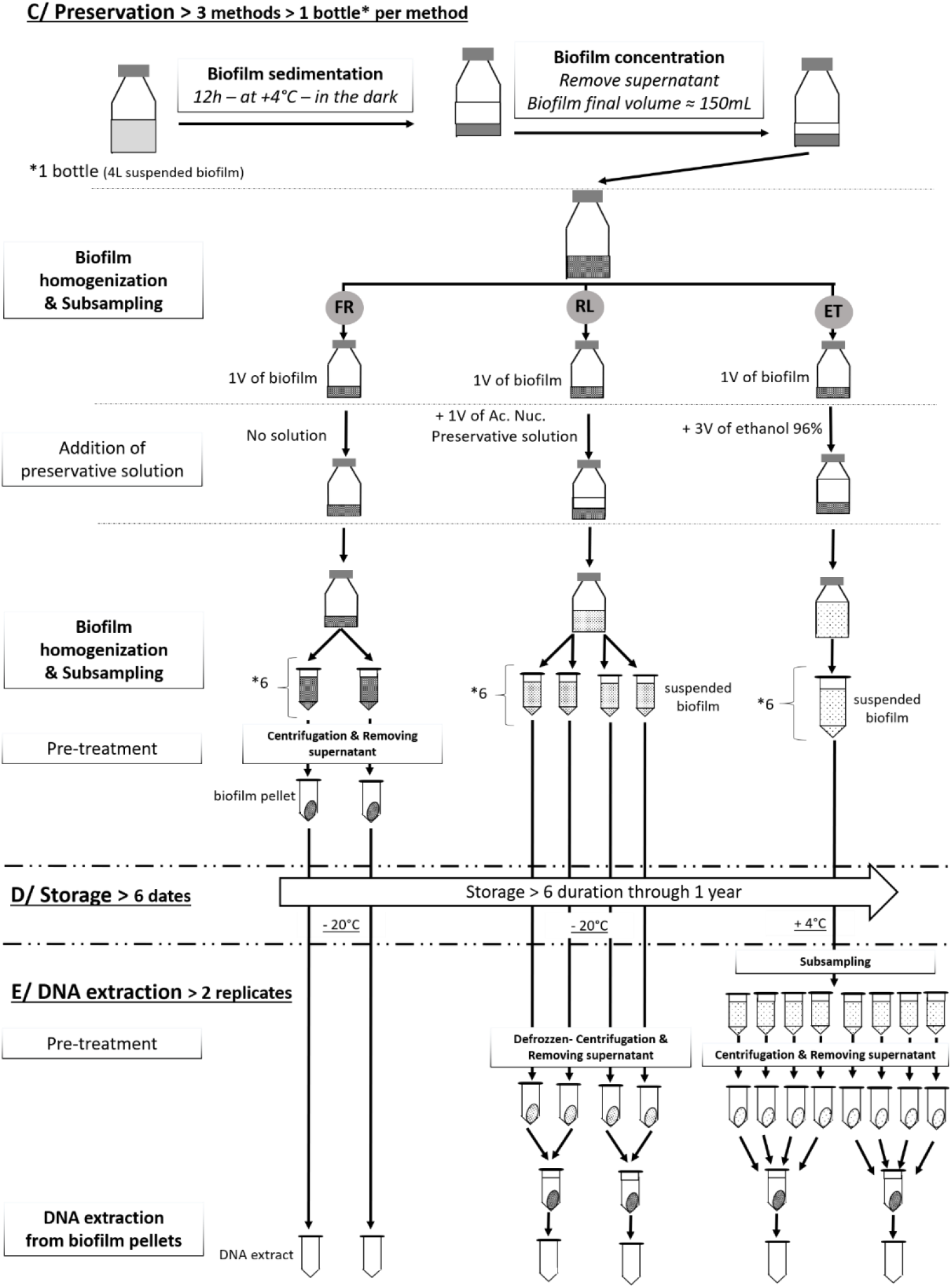
Detailed workflow presenting the three preservation methods (FR: deep-frozen, RL (nucleic acid preservative solution), and ET (ethanol)) for the freshwater biofilm samples. See Material and Methods for detailed explanations.

**Supp. Data 2.**
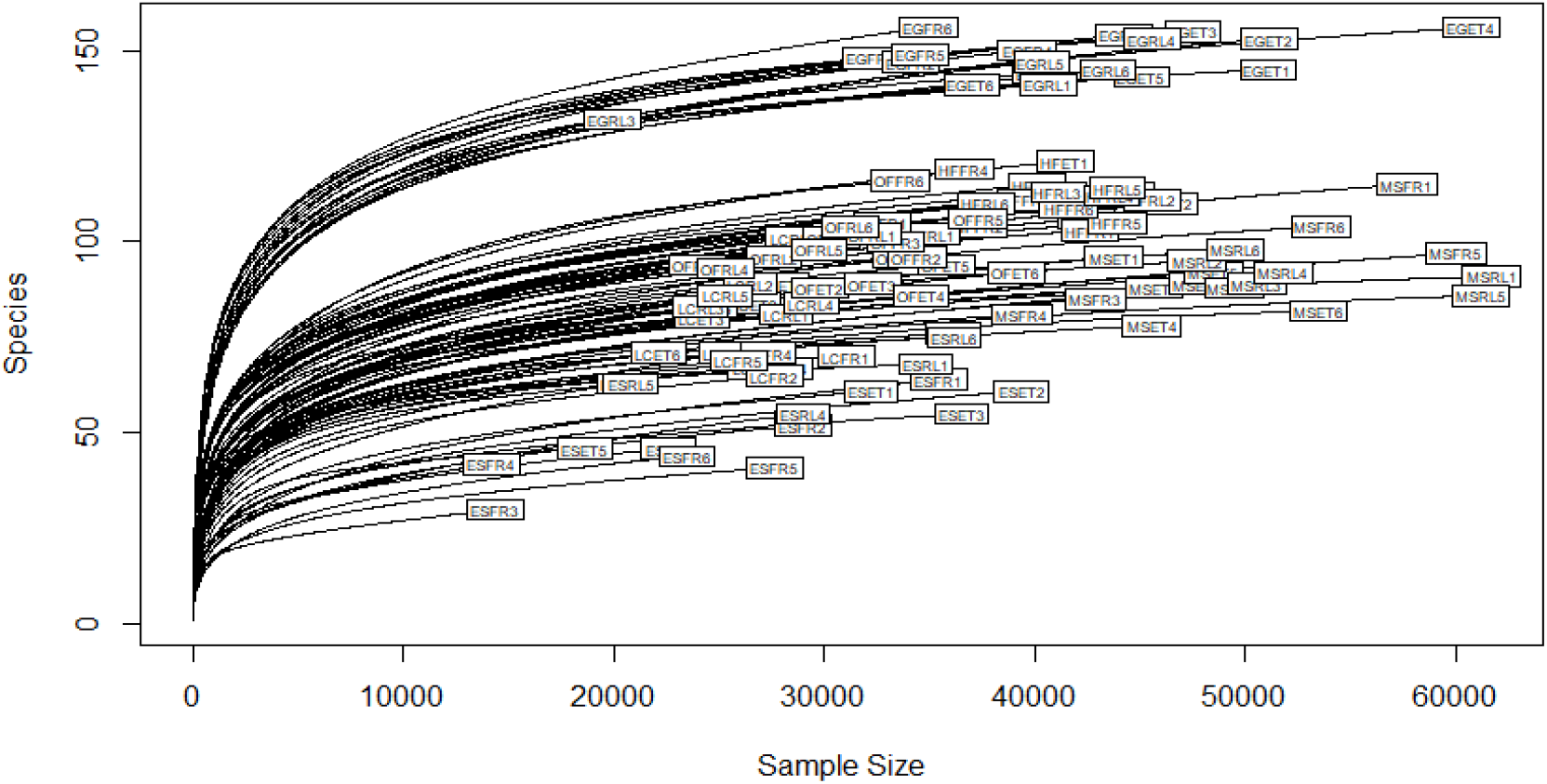
Sample rarefaction curves. Sample names are marked as a combination of codes for site, method and time.

**Supp. Data 3.**
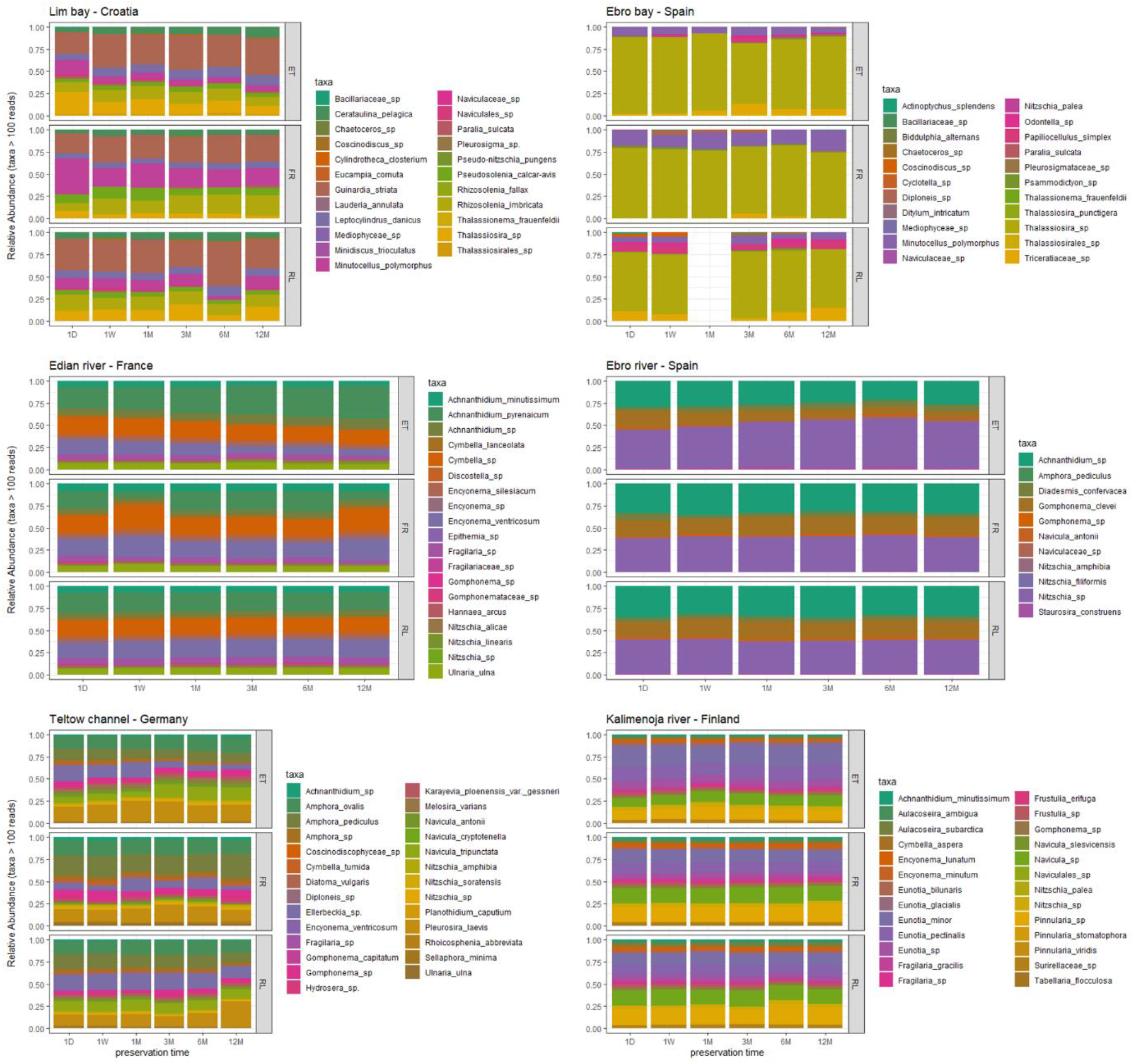
Diatom community composition assessed by DNA metabarcoding for marine sites (top row) and freshwater sites (middle and bottom rows) according to preservation methods (ET, FR, RL) and preservation duration (x axis). Only abundant taxa with more than 100 reads are shown.

